# Inconsistencies between human and macaque lesion data can be resolved with a stimulus-computable model of the ventral visual stream

**DOI:** 10.1101/2022.09.12.507636

**Authors:** Tyler Bonnen, Mark A.G. Eldridge

**Affiliations:** Stanford University, Stanford CA; Laboratory of Neuropsychology, National Institute of Mental Health, National Institutes of Health, Bethesda, MD

## Abstract

Decades of neuroscientific research has sought to understand medial temporal lobe (MTL) involvement in perception. The field has historically relied on *qualitative* accounts of perceptual processing (e.g. descriptions of stimuli), in order to interpret evidence across subjects, experiments, and species. Here we use *stimulus computable* methods to formalize MTL-dependent visual behaviors. We draw from a series of experiments (Eldridge et al., 2018) administered to monkeys with bilateral lesions that include perirhinal cortex (PRC), an MTL structure implicated in visual object perception. These stimuli were designed to maximize a *qualitative* perceptual property (‘feature ambiguity’) considered relevant to PRC function. We formalize perceptual demands imposed by these stimuli using a computational proxy for the primate ventral visual stream (VVS). When presented with the same images administered to experimental subjects, this VVS model predicts both PRC-intact and -lesioned choice behaviors; a linear readout of the VVS should be sufficient for performance on these tasks. Given the absence of PRC-related deficits on these ‘ambiguous’ stimuli, we (Eldridge et al., 2018) originally concluded that PRC is not involved in perception. Here we (Bonnen & Eldridge) reevaluate this claim. By situating these data alongside computational results from multiple studies administered to humans with naturally occurring PRC lesions, this work offers the first formal, cross-species evaluation of MTL involvement in perception. In doing so, we contribute to a growing understanding of visual processing that depends on—and is independent of—the MTL.

## 1 Introduction

Neuroanatomical structures within the medial temporal lobe (MTL) are known to support memory-related behaviors (Eichenbaum and Cohen, 2004; LaRocque and Wagner, 2015; Scoville and Milner, 1957). For decades, experimentalists have also observed MTL-related impairments in tasks designed to test perceptual processing (Baxter, 2009; Suzuki, 2009). In the object perception literature, these findings centered on perirhinal cortex (PRC), an MTL structure situated at the apex of high-level sensory cortices (Fig. 1a). Many studies reported visual impairments following lesions to PRC in humans and other animals, bolstering a perceptual-mnemonic account of perirhinal function (e.g. Murray and Bussey, 1999; Barense et al., 2007; Bussey et al., 2002; Inhoff et al., 2019; Lee et al., 2006; Lee et al., 2005). However, there were many visual experiments for which no PRC-related impairments were observed (e.g. Buffalo et al., 1998a; Buffalo et al., 1998b; Knutson et al., 2012; Stark and Squire, 2000). Decades of evidence resulted in a pattern of seemingly inconsistent experimental outcomes, with no formal method for disambiguating between competing interpretations of the available data.

**Figure 1:**
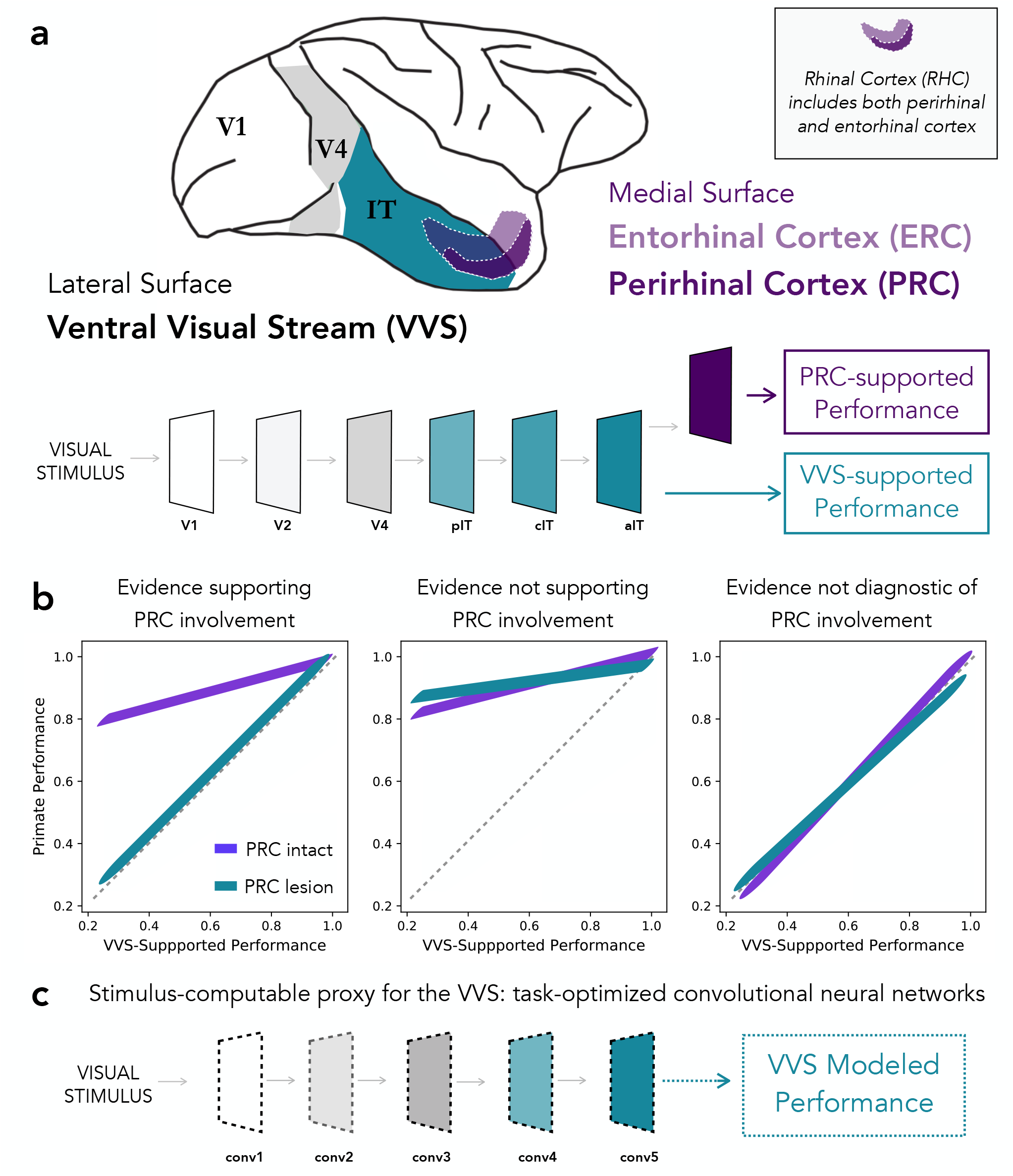
Formalizing MTL involvement in visual object perception. **(a)** Perirhinal cortex (PRC) is a poly-sensory medial temporal lobe structure situated at the apex of the primate ventral visual stream (VVS). Additionally, PRC lies within rhinal cortex (RHC; see inset). A perceptual-mnemonic hypothesis of PRC suggests that, in addition to its mnemonic functions, PRC enables ‘complex’ perceptual representations—i.e. those not supported by canonical sensory cortices alone. As such, PRC-related perceptual deficits are only expected for stimuli that require sufficiently ‘complex’ representations. PRC-intact subjects can outperform a direct readout from high-level visual cortex, and PRC-lesioned performance can be approximated by a computational model of the VVS (Bonnen et al., 2021). **(b)** We schematize the pattern of results that would be consistent with the PRC involvement in perception (left), results which indicate that non-PRC brain structures are required to outperform the VVS (center), as well as results which indicate that a visual discrimination task is supported by the VVS (i.e. no extra-VVS perceptual processing is required). **(c)** In order to evaluate VVS-supported performance on any arbitrary stimulus set, even in the absence of direct neural recordings, we leverage a computational model able to make predictions about VVS-supported performance, directly from experimental stimuli: task optimized convolutional neural networks.

A central challenge in this literature has been isolating PRC-dependent behaviors from those supported by PRC-adjacent sensory cortex. In the primate, this requires disentangling PRC-dependent performance from visual behaviors that depend on the ventral visual stream (VVS; DiCarlo and Cox, 2007; DiCarlo et al., 2012). Lacking more objective metrics, experimentalists had relied on informal, descriptive accounts of perceptual demands—terms such as ‘complexity’ and ‘feature ambiguity’—intended to characterize the stimulus properties that are necessary to evaluate PRC involvement in visual object perception. However, this informal approach enabled conflicting interpretations of the available evidence, without any means to arbitrate between them. For example, the absence of PRC-related deficits in a given study (e.g. Stark and Squire, 2000) has led some to conclude that PRC is not involved in perception (Suzuki, 2009), while others argue that stimuli in these studies can be represented by canonical visual cortices, and so no perceptual deficits are expected (Bussey and Saksida, 2002).

Recent work has provided a framework for assessing the perceptual demands of stimuli used to evaluate PRC involvement in visual object perception (Bonnen et al., 2021). Using a computational proxy for the VVS, Bonnen et al. analyzed visual discrimination experiments previously administered to humans with naturally occurring PRC lesions. The authors observed a striking correspondence between PRC-lesioned performance and a computational proxy for the VVS. Humans with an intact PRC, however, were able to outperform both PRC-lesioned subjects and VVS-modeled accuracy, as well as a linear readout of electrophysiological recordings from high-level visual cortex (schematized in Fig. 1d: left). This approach also identified numerous experiments in the literature that were not suitable for evaluating PRC contributions to perception; a computational proxy for the VVS achieved ceiling performance on these stimuli, suggesting no requirement for extra-VVS perceptual processing. This study resolved decades of seemingly inconsistent findings in the experimental literature, and the resulting data implicated PRC in visual object processing. Nonetheless, these analyses were restricted to just a sub-space of an experimental literature that spans many species and experimental designs.

Here we reanalyze data from visual classification tasks administered to PRC-lesioned and -intact monkeys (*macaca mulatta*), previously collected by Eldridge et al., 2018. Although we refer to these experimental groups as PRC intact/lesioned subjects, we note that lesioned subjects had bilateral damage to rhinal cortex (RHC), which includes both perirhinal and entorhinal cortex (Fig. 1a). We note that prior observation from Bonnen et al., 2021 unambiguously constrains how evidence from Eldridge et al., 2018 can be interpreted. If PRC-intact accuracy exceeds VVS-supported performance, it may be due to PRC-dependent contributions (schematized in Fig. 1b: left), or for reasons unrelated to PRC function (schematized in Fig. 1b: middle). However, if VVS-supported performance approximates PRC-intact behavior, no perceptual processing beyond the VVS should be necessary (schematized in Fig. 1b: right). We refer to the latter as ‘non-diagnostic’ stimuli. Using a computational proxy for the primate visual system, here we estimate VVS-supported performance on the categorization of stimuli from Eldridge et al., 2018, and compare model performance to PRC-intact and -lesioned macaque choice behaviors. We interpret these results in light of computational analysis of human participants with naturally occurring PRC lesions, enabling us to compare human and macaque lesion studies within a shared metric space (i.e. VVS model performance).

## 2 Results

We first extract model responses to each stimulus in all four experiments administered by Eldridge et al., 2018. In these experiments, subjects provided a binary classification for each stimulus: ‘cat’ or ‘dog.’ Critically, stimuli were composed not only of cats and dogs, but of ‘morphed’ images that parametrically vary the percent of category-relevant information present in each trial. For example, ‘10% morphs’ were 90% cat and 10% dog. These morphed stimuli were designed to evaluate PRC involvement in perception by creating maximal ‘feature ambiguity,’ a perceptual quality reported to elicit PRC dependence in previous work (Bussey et al., 2002, 2006; Murray and Richmond, 2001; Norman and Eacott, 2004). On each trial, subjects were rewarded for responses that correctly identify which category best fits the image presented (e.g. 10%=‘cat’, 90%=‘dog’, correct response is ‘dog’). We evaluate data from two groups of monkeys in this study: an unoperated control group (n=3) and a group with bilateral removal of rhinal cortex, which including peri- and ento-rhinal cortex. We formulate the modeling problem as a binary forced choice (i.e. ‘dog’=1, ‘cat’=0) and present the model with experimental stimuli. We then extract model responses from a layer that corresponds to ‘high-level’ visual cortex and learn a linear mapping from model responses to predict the category label. For all analyses, we report the results on held-out data (Methods: Determining model performance).

We first evaluate model performance with the aggregate metrics used by the original authors. With the original behavioral data we average performance across images within each morph level (e.g. 10%, 20%, etc.) across subjects in each lesion group (PRC-intact Fig. 2a, and -lesioned Fig. 2b). As reported in Eldridge et al., 2018, there is not a significant difference between the choice behaviors of PRC-lesioned and -intact subjects (no significant difference between PRC-intact/-lesion groups: *R*^2^ = 0.00, *β* = 0.01, *F* (1, 86) = 0.07, *P* = 0.941). For each of these experiments, we extract model responses to all stimuli from a model layer that best corresponds to a high-level visual region, inferior temporal (IT) cortex. Using the model responses from this ‘IT-like’ model layer to each image, we train a linear, binary classification model on the category label of each image (i.e. ‘dog’ or ‘cat’) on 4/5th of the available stimuli. We then evaluate model performance on the remaining 1/5th of those stimuli, repeating this procedure across 50 iterations of randomized train-test splits. A computational proxy for the VVS exhibits the same qualitative pattern of behavior as each subject group (Fig. 2c, model performance across multiple train-test iterations in black). Moreover, we observe a striking correspondence between model and PRC-intact behavior (Fig. 2d; purple: *R*^2^ = 0.98, *β* = 0.97, *t*(21) = 33.12, *P* = 6 × 10^*−*19^) as well as -lesioned subjects (green: *R*^2^ = 0.99, *β* = 0.96, *t*(21) = 57.38, *P* = 1 × 10^*−*23^). Employing the same metric used to claim no significant difference between PRC-lesion/-intact performance, we find no difference between subject and model behavior (*R*^2^ = 0.00, *β* = *−*0.01, *F* (1, 86) = *−*0.11, *P* = 0.915).

**Figure 2:**
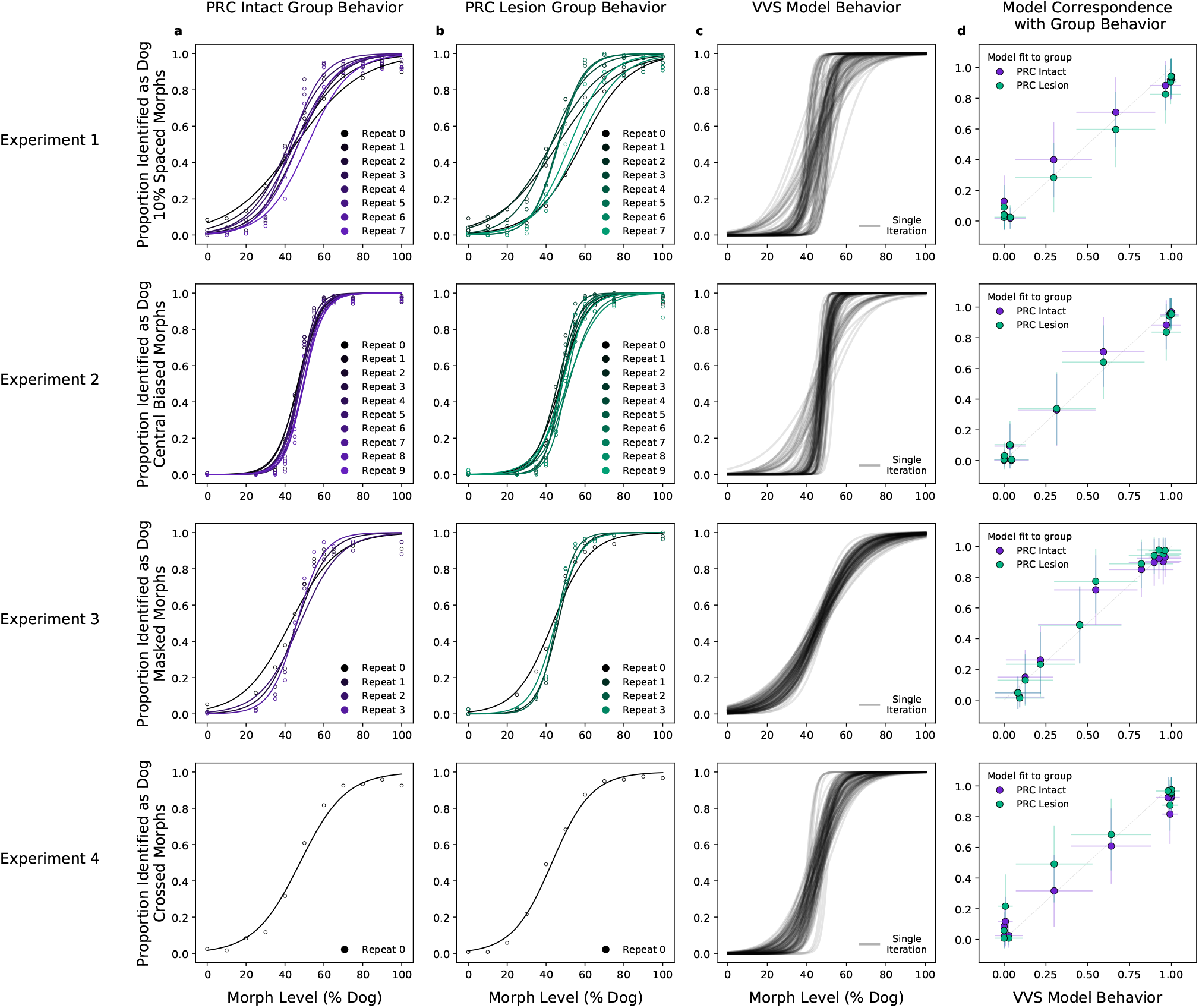
A computational proxy for the VVS predicts PRC-intact and -lesioned behavior. Averaging across subjects and morph-levels (i.e. all 10% morphs, 20% morphs, etc.), **(a)** PRC-intact (n=3) and **(b)** PRC-lesioned (n=3) subjects exhibit a similar pattern of responses across experiments (rows 1-4). We present stimuli used in this experiment to a computational proxy for the ventral visual stream (VVS), extracting model responses from a layer that corresponds with ‘high-level’ perceptual cortex. From these model responses we learn to predict the category membership of each stimulus, **(c)** testing this linear mapping on left-out images across multiple train-test iterations (black). **(d)** This computational proxy for the VVS accurately predicts the choice behavior of PRC-intact (purple) and -lesioned (green) grouped subjects (error bars indicate standard deviation from the mean, across model iterations and subject choice behaviors). As such, a linear readout of the VVS appears to be sufficient to perform these tasks, thus there need be no involvement of PRC to achieve neurotypical performance.

We extend our analysis beyond the aggregate morph- and subject-level analyses used by the original authors, introducing a split-half reliability analysis (Methods: Split-half reliability estimates). This enables us to determine if there is reliable choice behavior, for each subject, at the level of individual images. We restrict our analyses to experiments with sufficient data, as this analysis requires multiple repetitions of each image; we exclude experiments three (‘Masked Morphs’) and four (‘Crossed Morphs’) due to insufficient repetitions (which can be seen in Fig. 2, rows 3-4). Across both remaining experiments, we find consistent image-level choice behaviors for subjects with an intact (e.g. median 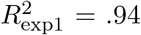, median 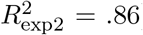) and lesioned (e.g. median 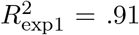, median 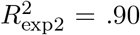) rhinal cortex (Fig. 3a: within subject reliability on the diagonal; PRC-intact subjects in purple, PRC-lesioned subjects in green). We also observe consistent image-level choice behaviors between subjects (e.g. median 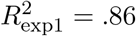, median 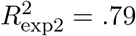). These results indicate there is reliable within- and between-subject variance in the image-by-image choice behaviors of experimental subjects (Fig. 3a: PRC-intact subjects purple, PRC-lesioned subjects green; between-group reliability in grey), suggesting that this behavior is a suitable target to evaluate how well we approximate more granular subject behaviors with a computational proxy for the VVS. We next examine whether the model can predict these more granular, image-level choice behaviors.

**Figure 3:**
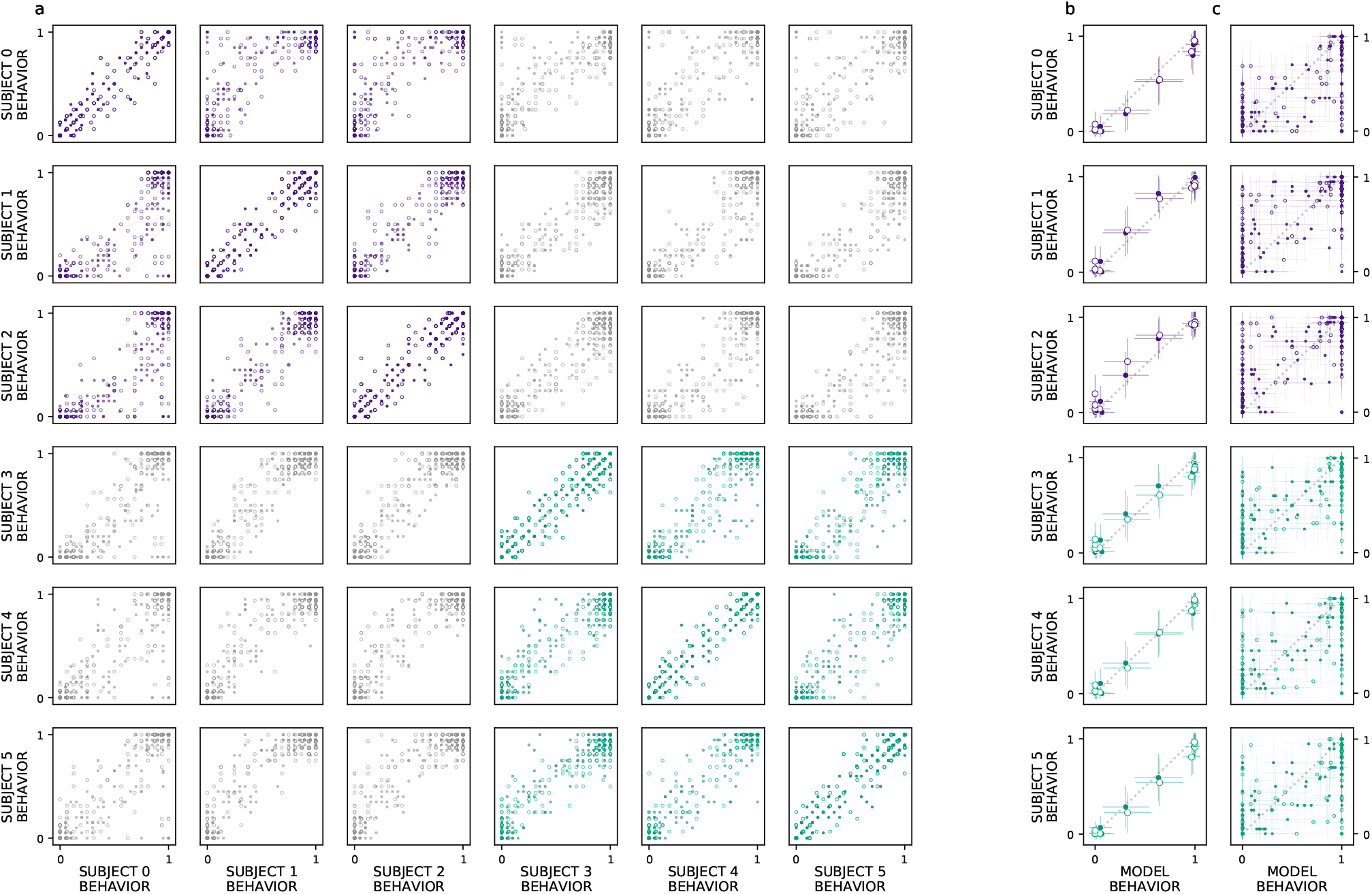
VVS model fits subject behavior for aggregate but not image-level metrics. Here we perform more granular analyses than those conducted by the authors of the original study: evaluating the model’s correspondence with PRC-lesioned and -intact performance at the level of individual subjects and images. We restrict ourselves to experiments that had sufficient data to determine the split-half reliability of each subject’s choice behaviors. First, we determine whether there is reliable image-level choice behavior observed for each subject, i.e. no longer averaged across morph levels. **(a)** We estimate the correspondence between subject choice behaviors over 100 split-half iterations, for both Experiments 1 (closed circles) and 2 (open circles), using *R*^2^ as a measure of fit. Each row contains a given subjects’ (e.g. subject 0, top row) correspondence with all other subjects’ choice behaviors, for PRC-intact (purple) and -lesioned (green) subjects. We find that the image-level choice behaviors are highly reliable both within- (on diagonal) and between-subject (off diagonal), including between PRC-lesioned and -intact subjects (grey). We next compare model performance to the behavior of individual subjects, averaging over morph levels in accordance with previous analyses (i.e. averaging performance across all images within each morph level, e.g. 10%). **(b)** We observe a striking correspondence between the model and both PRC-lesioned (green) and PRC-intact (purple) performance for all subjects. **(c)** Finally, for each subject, we estimate the correspondence between model performance and the subject-level choice behaviors, at the resolution of individual images. Although model fits to subject behavior are statistically significant, it clearly does not exhibit ‘subject-like’ choice behavior at this resolution.

Our computational approach is able to predict subject-level choice behavior when aggregated across morph levels, for both PRC-intact (e.g. subject 0; *R*^2^ = 0.99 *β* = 1.01, *t*(21) = 39.30, *P* = 2 × 10^*−*20^) and -lesioned (e.g. subject 4: *R*^2^ = 0.99 *β* = 1.01, *t*(21) = 45.01, *P* = 1 × 10^*−*21^) subjects (Fig. 3b). Interestingly, the model’s fit to subject behavior is indistinguishable from the distribution of between-subject reliability estimates (Fig. S4a; median of the empirical P(model|reliability_between-subject_) = .592) suggesting that the model exhibits ‘subject-like’ behaviors at this resolution. Our modeling approach is also able to significantly predict image-level choice behaviors for both PRC-lesioned (e.g. subject 3: *R*^2^ = 0.86 *β* = 0.81, *F* (1, 438) = 52.79, *P* = 5 × 10^*−*192^) and -intact subjects (e.g. subject 1: *R*^2^ = 0.87 *β* = 0.88, *F* (1, 438) = 53.24, *P* = 2 × 10^*−*193^). However, the model behavior is unlikely to be observed under the distribution of between-subject reliability estimates (between-subject reliability distributions visualized in Fig. S4b; median of the empirical P(model|reliability_between-subject_) = 0). That is, the model does not exhibit ‘subject-like’ choice behaviors at the resolution of individual images—as previously reported (Rajalingham et al., 2018).

There are properties of the experimental design in Eldridge et al., 2018 that encourage a more careful comparison between primate and model behavior. Experimental stimuli contain discrete interpolations between ‘cat’ and ‘dog’ images, such that adjacent stimuli within a morph sequence are highly similar (for examples see Supplemental Fig. S5). The co-linearity in this stimulus set is revealed by running a classification analysis over pixels: a linear readout of stimulus category directly from the vectorized (i.e. flattened) images themselves is sufficient to approximate aggregate performance of all experimental groups (*R*^2^ = 0.94 *β* = 0.90, *F* (1, 42) = 26.74, *P* = 5 × 10^*−*28^; Supplemental Fig. S6). To ensure that the VVS modeling approach is not simply a byproduct of the co-linearity in the stimuli, we construct a conservative method for model evaluation by restricting training data to images from unrelated morphs sequences (i.e. train on morph sequences A-F, test on morph sequence G). Under this more conservative train-test split, pixels are no longer predictive of primate behavior (*R*^2^ = 0.05 *β* = 0.45, *F* (1, 42) = 1.42, *P* = 0.164; Supplemental Fig. S7a-c), but there remains a clear correspondence between the model and PRC-lesioned (*R*^2^ = 0.87 *β* = 1.17, *F* (1, 42) = 16.94, *P* = 2 × 10^*−*20^) and -intact performance (*R*^2^ = 0.88 *β* = 1.16, *F* (1, 42) = 17.39, *P* = 8 × 10^*−*21^; Supplemental Fig. S8). That is, although subjects were able to exploit the colinearity in the stimuli to improve their performance with experience, the correspondence between VVS-models and primate choice behaviors is not an artifact of these low-level stimulus attributes.

## 3 Discussion

To evaluate competing claims surrounding PRC involvement in perception, Eldridge et al., 2018 administered a series of visual classification tasks to PRC-lesioned/-intact monkeys. These stimuli were carefully crafted to exhibit a qualitative, perceptual property that had previously been shown to elicit PRC dependence (i.e. ‘feature ambiguity’; Bussey et al., 2002, 2006; Murray and Richmond, 2001; Norman and Eacott, 2004). The absence of PRC-related deficits across four experiments led the original authors to suggest that perceptual processing is not dependent on PRC. Here we reevaluate this claim by situating these results within a more formal computational framework; leveraging task-optimized convolutional neural networks as a proxy for primate visual processing (Rajalingham et al., 2018; Schrimpf et al., 2020; Yamins et al., 2014). We first determined VVS-model performance on the experimental stimuli in Eldridge et al., 2018. We then compared these computational results with monkey choice behaviors, including subjects with bilateral lesions to PRC (n=3), as well as un-operated controls (n=3). For both PRC-lesioned/-intact monkeys, we observe a striking correspondence between VVS-model and experimental behavior at the group (Fig. 2d) and subject level (Fig. 3b). These results suggest that a linear readout of the VVS should be sufficient to enable the visual classification behaviors in Eldridge et al., 2018; no PRC-related impairments are expected.

In isolation, it is ambiguous how these data should be interpreted. For example, if VVS-modeled accuracy was sufficient to explain PRC-intact performance across all known stimulus sets, this would suggest that PRC is not involved in visual object perception. However, previous computational results from humans demonstrate that PRC-intact participants are able to outperform a linear readout of the VVS (schematized in Fig. 1b: left). Because results from these human experiments are in the same metric space as our current results (i.e. VVS-modeled performance), these data unambiguously constrain our interpretation: for a stimulus set to evaluate PRC involvement in visual processing, participants must be able to outperform a linear readout of the VVS. That is, supra-VVS performance must be observed in order to isolate PRC contributions from those of other possible contributors to these behaviors (e.g. prefrontal cortex, schematized in Fig. 1b: center). Given that supra-VVS performance is not observed in the current stimulus set (Fig. 2d; schematized in Fig. 1b: right), we conclude that experiments in Eldridge et al., 2018 are not diagnostic of PRC involvement in perception. Consequently, we suggest that these data do not offer absolute evidence against PRC involvement in perception— revising the original conclusions made from this study.

We note that there is meaningful variance in the primate behavioral data not captured by the current modeling framework. The original analyses in Eldridge et al., 2018 were performed at the ‘morph’ level (i.e. averaging across multiple images within the same ‘morph’ percent; Supplementary Fig. S5); at this coarse resolution of behavior, the model exhibits subject-like choice performance (Supplemental Fig. S4a). In addition to these original metrics, here we conducted a more granular analyses at the level of individual images (i.e. averaging across multiple presentations of the same image, not across multiple images within the same morph level). We find that, for PRC-lesioned and -intact monkeys, choice behaviors are reliable, both within- and between-subjects (Fig. 3a). At this image-level resolution, however, the VVS model does not match the pattern of choice behaviors from experimental subjects (Fig. 3c; Supplemental Fig. S4b). This observation is consistent with previous reports (Rajalingham et al., 2018), suggesting that these VVS-like models are best suited to approximate aggregate choice behaviors, not responses to individual images. Many sources of variance have been identified as possible contributors to subject-model divergences, such as their deterministic, strictly feedforward processing (Kar and DiCarlo, 2020). Nonetheless, these admittedly coarse models of visual information processing provide us with leverage to formalize theories of MTL involvement in visual discrimination behaviors. Although stimuli in Eldridge et al., 2018 do not elicit PRC involvement, this computational framework has revealed PRC dependent visual processing in other studies (Bonnen et al., 2021). What stimulus attributes require visual processing by PRC? Many *descriptive* accounts have been offered, encapsulated by terms such as ‘feature ambiguity’ (Bussey and Saksida, 2002), stimulus ‘complexity’ (Murray and Bussey, 1999), and ‘configural’ processing (Barense et al., 2007). While operationalizing these terms across studies has proven challenging, several heuristics have emerged. First, viewpoint variation appears to play a critical role in eliciting PRC involvement, as has previously been suggested (Barense et al., 2010). Despite stimuli in Eldridge et al., 2018 being designed to maximize ‘feature overlap,’ these stimuli have no viewpoint variation; the absence of PRC-related deficits is consistent with the interpretation that variation in viewpoint might be necessary for recruiting PRC. Second, PRC-related deficits appear to be especially pronounced for ‘unfamiliar’ stimuli (Barense et al., 2007, Liang et al., 2020), though this can be challenging concept to parametrize. While these heuristics (e.g. viewpoint variation and perceptual novelty) inform our understanding of MTL-dependent visual processing, our results showcase the need to incorporate stimulus computable methods when designing novel stimuli and evaluating experimental evidence.

Which experimental designs are best suited to evaluate PRC-involvement in perception? Historically, neuroscientists have sought to isolate PRC-related visual processing using task designs that minimize memory-related demands. This has led to the widespread use of visual discrimination tasks that present all stimuli on screen concurrently (‘oddity’ tasks, e.g. Barense et al., 2007; Bussey and Saksida, 2002) as well as classification (Eldridge et al., 2018) and binary forced choice (Bussey et al., 2002) experiments. Each of these designs provides unique insights, but synthesizing these results has proven challenging. Fortunately, the current modeling framework is agnostic to many of these design choices; application of these computational tools can bring experimental results across these experimental settings into a unified metric space (i.e. model performance) enabling a formal comparison of these data across studies and across species. Finally, we note that there are temporal dynamics which are characteristic of PRC-dependent visual processing across multiple experimental designs: the magnitude of divergence between PRC- and VVS-supported performance scales, linearly, with reaction time (Bonnen et al., 2021), presumably in order to enable subjects to collect multiple visual samples via eye movements (Erez et al., 2013). We believe that these computational tools, alongside a careful consideration of animal behavior, will enrich the next generation of empirical studies surrounding MTL-dependent perceptual processing.

## 4 Acknowledgements

This work is supported by the Intramural Research Program, National Institute of Mental Health, National Institutes of Health, Department of Health and Human Services (annual report number ZI-AMH002032) as well as the NIH Blueprint D-SPAN Award (F99/K00; 1F99NS125816-01) and Stanford’s Center for Mind Brain Behavior and Technology. We thank Elizabeth Murray and Anthony Wagner for insightful conversations and suggestions on this manuscript and throughout the course of this work.

## 5 Methods

### 5.1 Determining model performance

For all estimates of model performance we use a task-optimized convolutional neural network pretrained on Imagenet (Deng et al., 2009). For transparency, we report the results from the first instance of this model class used to evaluate these data (Simonyan and Zisserman, 2014), but note that these results hold across all model instances evaluated. We preprocess each image from Eldridge et al., 2018 using a standard computer vision preprocessing pipeline; resizing images to a width and height of 224×224, then normalizing each image by the mean ([0.485, 0.456, 0.406]) and standard deviation ([0.229, 0.224, 0.225]) of the distribution of images used to train this model. We extract model responses to each image from a layer (fc6) previously shown to exhibit a high correspondence with electrophysiological responses to high-level visual cortex (Bonnen et al., 2021). We use a l2-normed logistic regression model implemented in sklearn (Pedregosa et al., 2011) to predict the binary category classification (i.e. ‘dog’=1, ‘cat’=0). For each experiment, we first estimate the optimal regularization strength (‘C’) for the logistic regression model through 5-fold cross validation (10^*−*5^ to 10^*−*5^) on 4/5th of the data. We then evaluate model performance on each experiment on independent data (i.e. the remaining 1/5th of stimuli).

### 5.2 Consistency estimates

We estimate within- and between-subject consistency using a common protocol. For the given resolution of analysis (either morph- or image-level), we require multiple presentations of the same items. For the morph-level analysis, which aggregates stimuli within ‘morph levels’ (e.g. all stimuli that are designed to be 0% dog, 10%, etc.), all stimulus sets meet this criterion. There are, however, multiple experiments that do not contain sufficient data to perform the image-level analysis, which requires multiple presentations of each stimulus; experiment four contains only one presentation of each stimulus, precluding it from our consistency analyses, and experiment three contains only 4 repetitions, which is insufficient for reliable within- and between-subject consistency estimates. Thus, we restrict our consistency estimates to experiments one (10 repetitions per image) and two (8 repetitions per image). All repetition counts are evident in Fig. 2.

We estimate all consistency metrics over 100 random split-halves iterations. For each iteration, across all items within a given resolution (where items can refer to either a given morph percent, for the morph-level analysis, or a given image, for the image-level analysis), we randomly split choice behavior into two random splits. In the image-level analysis, for example, for each image *x*_*i*_ within the set of *n* images, we randomly select half of all repetitions of 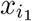, and compute the mean of this random sample 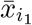, for all *n* images. We repeat this procedure for the remaining half, 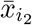, across all *n* images. Thus, we have two *n* dimensional vectors, 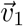 and 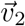, randomly sampled from the full distribution of choice behaviors. We use *R*^2^ as a measure of fit between each split half. We repeat this measure over 100 iterations, resulting in a distribution of split-half fits. For within-subject consistency metrics, we generate each split by randomly sampling from a single subjects’ choice behaviors. For the between subject consistency metrics, we compute 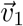 from subject_*i*_s choice behavior, and 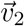 from subject_*j*_s choice behavior, using the same protocol described above.

## 6 Supplemental Information

**Figure S4:**
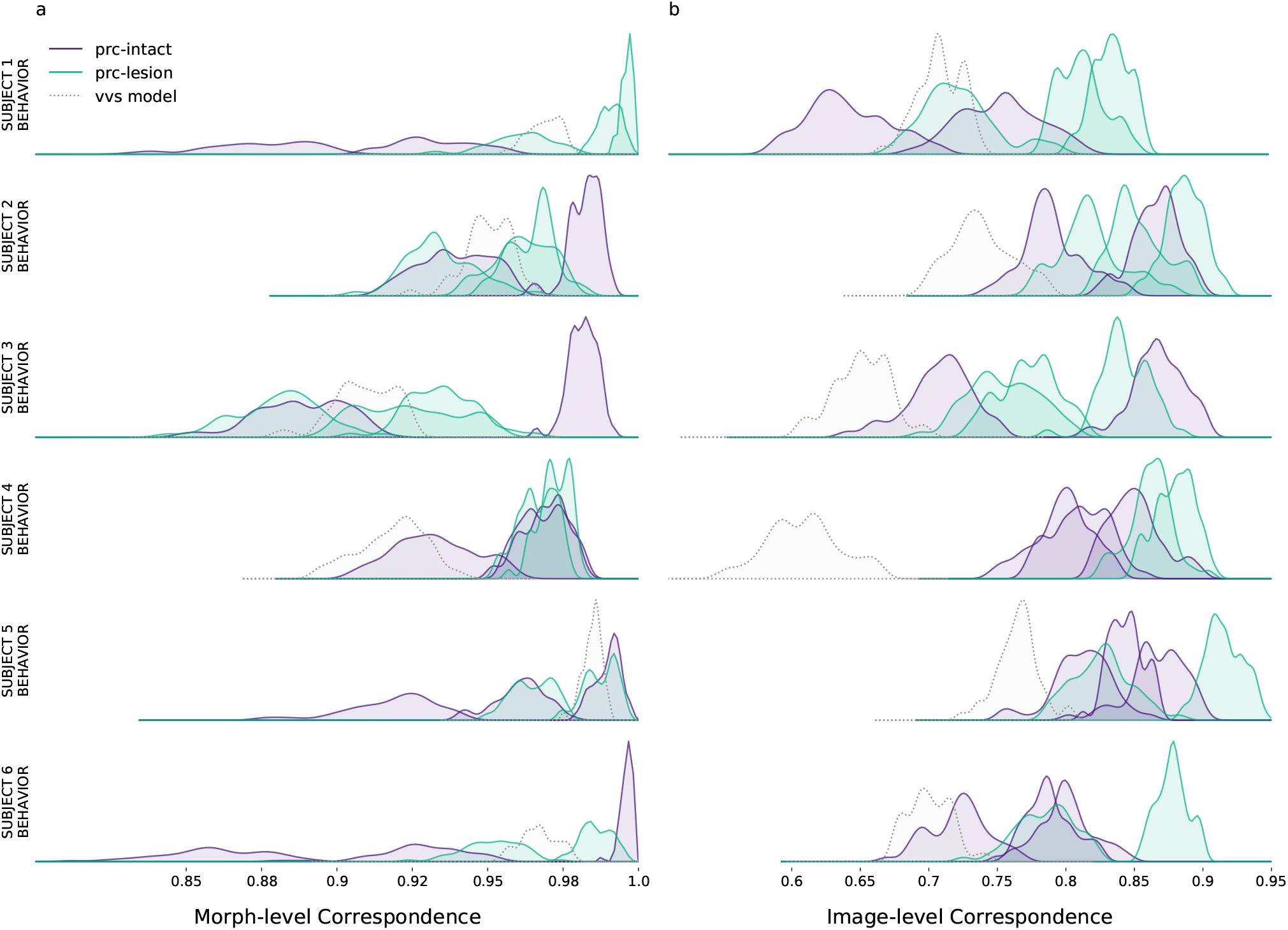
VVS model is ‘subject-like’ for aggregate but not image-level metrics. We estimate the correspondence between subject-subject choice behaviors. First, we generate a random split of each subjects performance. We then compute the between-subject correlation, iterating across 100 random splits. Each row contains a given subjects’ (e.g. subject 0, top row) correspondence with all other experimental subjects, including PRC-intact (purple) and -lesioned (green) monkeys. Using this same subject-subject measure, we also estimate subject-model correspondence (grey). We visualize our results at two resolutions: **(a)** for the morph-level analysis, we average performance across all images within each morph level (e.g. 10%, 20%, etc.; as per the analysis in Fig. 3b) and compare a single subject’s behaviors to all other experimental subjects, as well as model performance; **(b)** for the image-level analysis we average performance across all repetitions of each image (as per the analyses in Fig. 3c) and compare a single subject’s behaviors to all other experimental subjects, as well as model performance. For the morph-level analysis, the model choice behavior is ‘subject-like’; the distribution of model-subject correspondence is within the distribution of between-subject correspondence (in Fig. 3b, subject-level choices behaviors are on the diagonal). However, at the resolution of single images, model choice behavior is not subject-like; model correspondence to each subject is not likely observed under the between-subject distributions (in Fig. 3c, subject-level choices behaviors do not fall along the diagonal). We note the PRC-intact monkeys are subjects 1-3, PRC-lesioned monkeys are subjects 4-6

**Figure S5:**
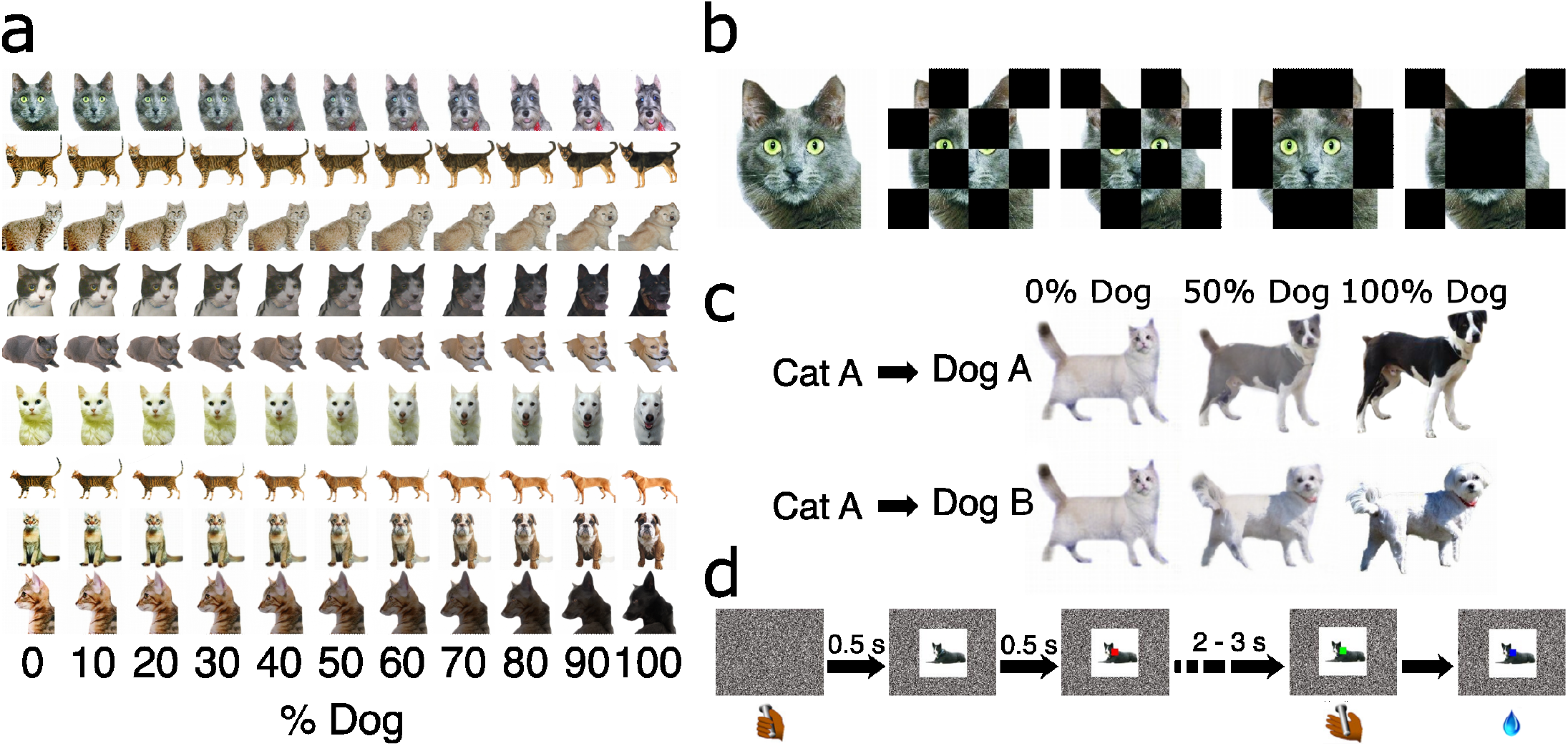
Experimental stimuli and protocol from Eldridge et al., 2018. **(a)** Example stimuli from experiment one, illustrating multiple instances of stimuli across morph levels. **(b)** Example stimuli used used for masked morphs, in experiment 3. **(c)** Example stimuli used for ‘crossed morphs’ in experiment 2. **(d)** Protocol for all experiments. Subject’s initiate each trial with a lever press. A stimulus is presented, followed by a red dot at the central field of view. Subjects could avoid an extended inter-trial delay by releasing the bar in the first interval (signaled by a red target) for stimuli that were less than 50% dog, and were rewarded for releasing the bar in the second interval (signaled by a green target) for stimuli that were more than 50% dog. This amounts to an asymmetrical reward structure. They were rewarded randomly for releasing during the green interval for 50 – 50 morphs.

**Figure S6:**
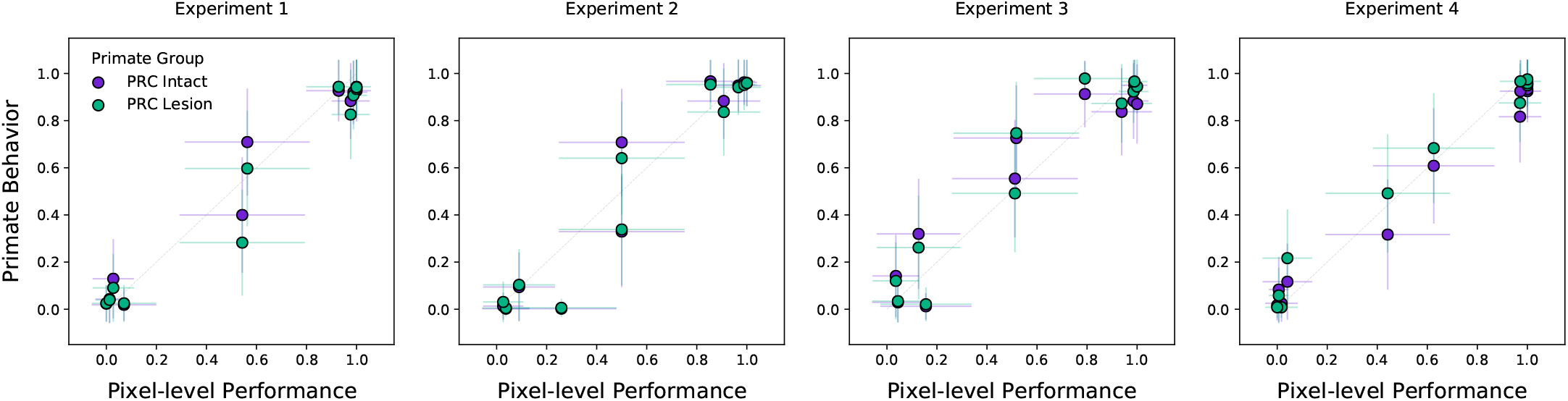
Colinearity within the stimulus set revealed by a pixel-level analysis. Classification behaviors on this stimulus set are learned and evaluated on images with a high degree of colinearity: stimuli with similar correct answers are highly overlapping, as can be seen in the example stimuli in Fig. S5. We formalize this observation by adapting previous analyses (visualized in Fig. 2d): In the place of a computational proxy for the primate VVS we use raw pixel-level representations, training a linear readout of stimulus category directly from the vectorized (i.e. flattened) images. We find that these pixel-level representations are sufficient to achieve the performance observed across experimental subjects in both PRC-intact and -lesioned groups. The co-linearity in these data suggest that ‘high-level’ representations may not be necessary to classify stimuli in these experiments, as a linear operation over the stimuli themselves achieve group-level performance.

**Figure S7:**
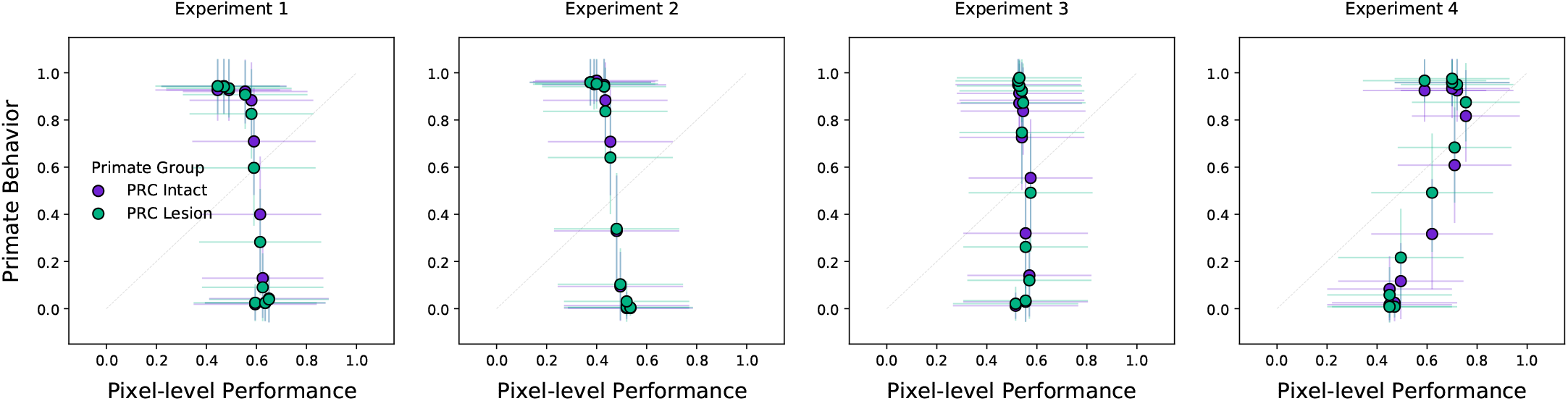
Pixel-level performance fails on a more conservative evaluation metric. Previous evaluation of model performance employed a train-test split that was ‘naive’ to the co-linearity in the available data. Here we implement and validate a more conservative evaluation metric: To evaluate each image within a given morph sequence (i.e. a unique cat-dog combination that spans from 0-100% dog) we remove all instances in that morph sequence from the training data. This ensures that there are no image-adjacent stimuli that the model can exploit (e.g. training on 10% and testing on 0% within the same morph sequence). As anticipated, pixel-level classification only achieves chance performance under this more conservative evaluation metric.

**Figure S8:**
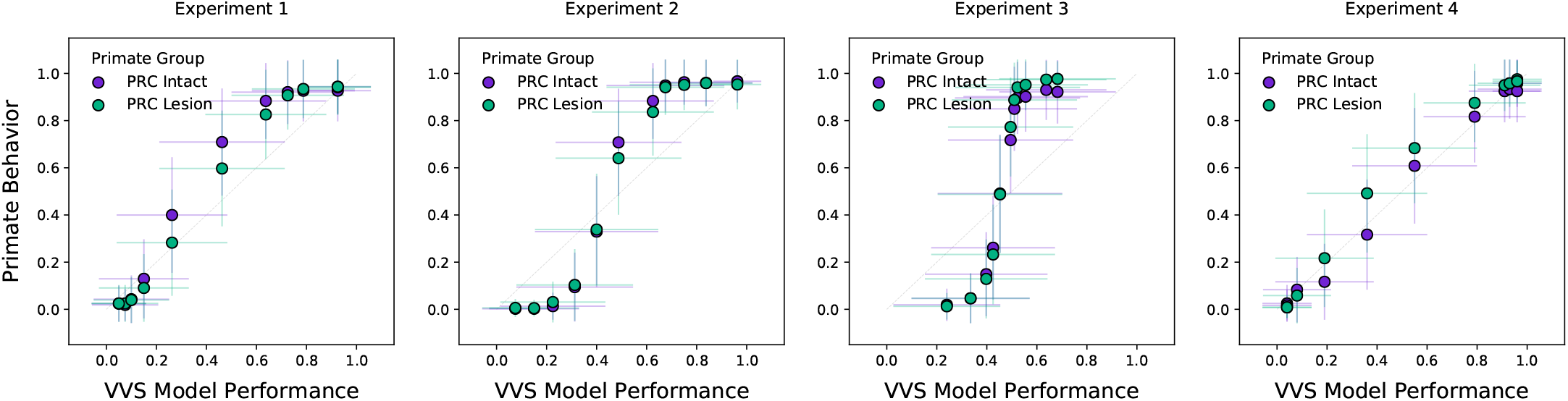
Model approximates primate behavior even with a more conservative evaluation metric. Here we evaluate model performance using a more conservative train-test split. While pixel-level representations in Fig. S7 fail to approximate group-level performance, a computational proxy for the VVS continues to predict group-level performance of both PRC-intact and -lesioned participants across experiments.

## Notes

### Competing Interest Statement

The authors have declared no competing interest.

